# READemption 2: Multi-species RNA-Seq made easy

**DOI:** 10.1101/2022.09.30.510338

**Authors:** Till Sauerwein, Thorsten Bischler, Konrad U. Förstner

## Abstract

Dual or Multi RNA-seq simultaneously analyze the transcriptomes of two or more interacting species to gain insights about their interplay. The RNA of the interacting species is collected and sequenced together and only separated *in silico* by mapping the reads to the corresponding genomes. We developed READemption 2.0, to our knowledge the first tool that performs all necessary steps to handle RNA-seq data from any number of species. These steps comprise basic quality filtering and adapter trimming of raw reads, aligning the reads to reference genomes, generating nucleotide-wise coverage files, creating gene-wise read counts and performing differential gene expression analysis. These results can be visualized by additional subcommands of the software. READemption 2.0 allows users to produce meaningful results with default settings that follow conventional standards. Furthermore, many parameters can be adjusted to meet the users’ specific needs, e.g. keeping or discarding species cross-mapped reads or normalizing the data.

## Introduction

Dual RNA-sequencing (Dual RNA-seq) is the simultaneous transcriptome profiling of two interacting species (***Westermann et al., 2012***). If more than two species are investigated the term Multi RNA-sequencing (Multi RNA-seq) is used. The distinctive feature of these methods, compared to conventional RNA-seq, is that the RNA of all interacting partners like a pathogen and its host is extracted and sequenced without physical separation. Since the RNA of different species is sequenced together, assigning each read to its originating species only happens *in silico* (Figure 1). The simultaneous investigation of two (or more) species allows researchers to correlate the transcriptome profiles and thus gain new insights of the molecular interplay of the interacting species.

**Figure 1.**
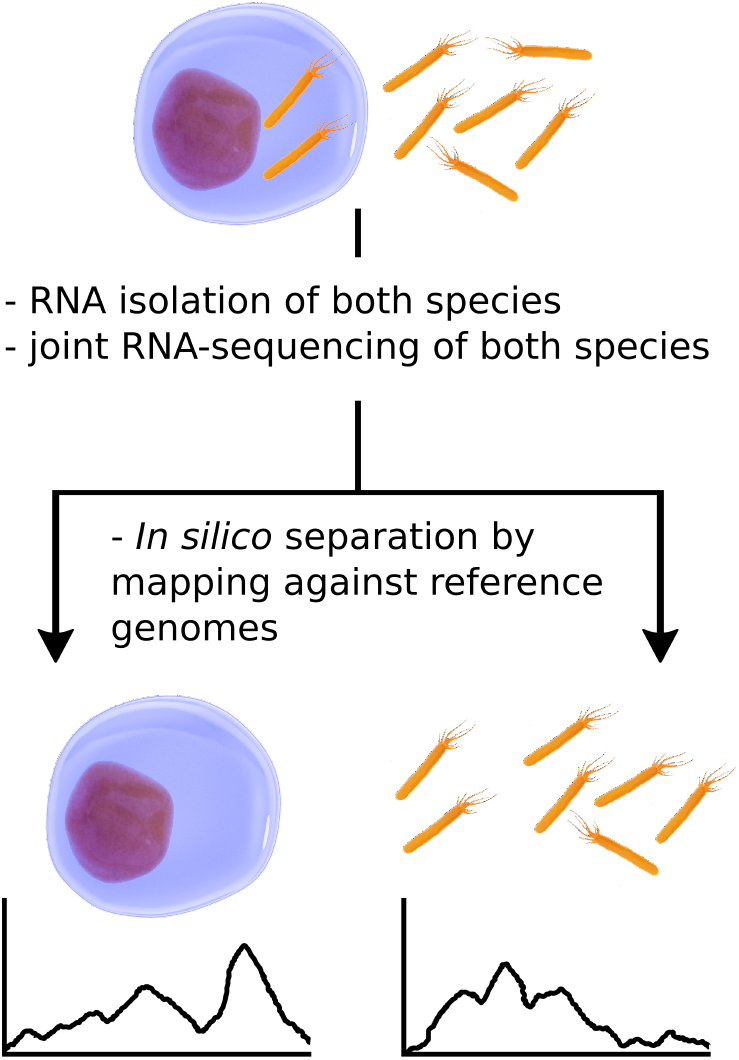
General Dual RNA-sequencing workflow.

Since the first application of Dual RNA-seq to an eukaryotic pathogen and host system (***Tierney et al., 2012***), and its theoretically assessment of the general feasibility in pathogen host systems in the early 2010s (***Westermann et al., 2012***), the method has been applied to a variety of hostpathogen, mutualistic and commensal interaction systems (***Wolf et al., 2018***).

Several recent studies investigated host-pathogen interactions: For example, ***Aulicino et al. (2022)*** revealed a dynamic adaption of iron metabolism during *Salmonella* infection of dendritic cells for both the human host and the bacterial pathogen. Different Salmonella strains used different evasive strategies to counteract the iron-driven antimicrobial defense of the host, which in turn showed unique responses depending on the infecting strain. *Staphylococcus aureus* showed differential expression of virulence factors during infection of two mice strains. The virulence was influenced by the host’s different level of resistance to the bacteria (***Thänert et al., 2017***). A recent study applied Dual RNA-seq to different SARS-CoV-2-infected patient samples and cell lines that revealed co-expressed viral and human genes. A consensus network derived from co-expression highlighted a host response characterized by increased chemokine and cytokine activity (***Maulding et al., 2022***).

Here we present READemption 2.0, an open source command line tool that allows users to analyze Dual or Multi RNA-seq data. To our knowledge, READemption 2.0 is the first tool that allows researchers to perform multi-species RNA-seq analysis with any number of species.

## Results

### Application and usage of READemption and the need for a Dual/Multi RNA-seq anal-ysis tool

Since READemptiorïs initial release in 2014 (***Förstner et al., 2014***) it has been used by numerous publications for analyzing data from different RNA-seq applications. Among these applications are conventional RNA-seq (***Aguilar et al., 2020; Lee et al., 2021***), differential RNA-seq (***Ponath et al., 2021; Ryan et al., 2020***), Grad-seq (***Hör et al., 2020; Smirnov et al., 2016***), RIP-seq (***Kavita et al., 2022; Liao et al., 2022***), CLIP-seq (***Bauriedl et al., 2020; Holmqvist et al., 2016***), TIER-seq (***Hoyos et al., 2020; Chao et al., 2017***) and metatranscriptomics (***Krohn-Molt et al., 2017***). The essential RNA-seq results that can be generated with READemption, like alignment files (BAM file format https://samtools.github.io/hts-specs/SAMv1.pdf), mapping statistics, nucleotide-wise coverage files, gene-wise quantification counts and differential gene expression analysis also serve as input for follow-up analysis tools: ANNOgesic (***Yu et al., 2018***), a tool for annotating bacterial and archaeal genomes uses coverage files as input e.g. for transcript start site and processing site detec-tion, sRNA (small RNA) detection and sRNA tar-get detection. GRADitude (https://github.com/foerstner-lab/GRADitude) uses gene-wise quantification counts and mapping statistics for RNA-RNA and RNA-protein prediction of GRAD-seq experiments. ClusterProfiler (***Wu et al., 2021***) performs gene set enrichment analysis (GSEA), which requires tables containing genes and their corresponding differential gene expression fold changes calculated by e.g. DESeq2 (***Love et al., 2014***), which is integrated into READemption’s subcommand ‘deseq’. PEAKachu (https://github.com/tbischler/PEAKachu), a peak-calling tool for CLIP-seq data, needs BAM files as input that can be generated with READemption’s ‘align’ subcommand.

The number of publications using READemption for RNA-seq analysis has increased over the years (Figure 2A, ***PubMed (2022a)***). This increase and the different RNA-seq protocols READemption has been applied to, show the need for an RNA-seq analysis tool that covers a broad spectrum of RNA-seq applications. As the number of publications applying Dual RNA-seq also increased over the years (Figure 2B, ***PubMed (2022b)***) and READemption could not handle Dual or Multi RNA-seq data without additional manual manipulation of input and output files, we developed READemption 2.0.

**Figure 2.**
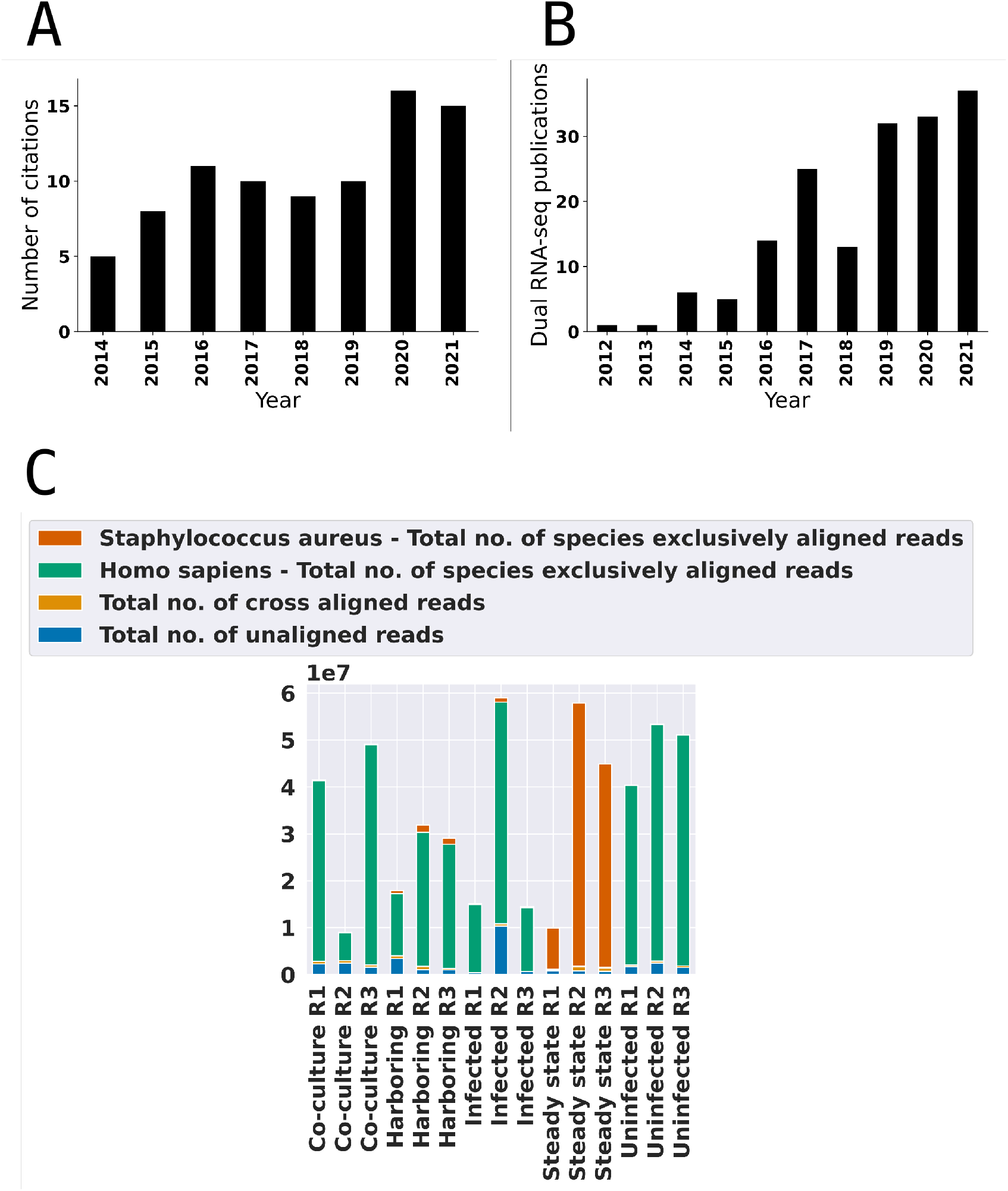
**(A)** Number of publications citing READemption 1.0 or earlier versions per year. **(B)** Number of publications having “Dual RNA-seq” in their title or abstract per year. **(C)** Alignment statistics plot of a Dual RNA-seq experiment with 15 libraries generated with READemption 2.0’s ‘viz_align’ subcommand. The plot shows the number of species exclusive aligned reads for each species, the species cross-mapped reads and the unaligned reads.

### READemption 2.0’s general workflow

READemption 2.0 is a major upgrade of the previous READemption RNA-seq analysis tool. The new version enables users to analyze projects with more than one interacting species, while keeping all of READemption 1.0’s core functionalities, including the ability to analyze RNA-seq data of a single species. To allow users an easy transition to the new version, the main workflow has not been changed and is as follows (see Figure 3): The first step of every new project is creating the input folder structure for reads, reference sequences and annotations with the subcommand ‘create’. READemption 2.0 adds the possibility to create annotation and reference sequence input folders for each species by providing individual names for each species being part of the current RNA-seq analysis project (the two species of the example workflow in Figure 3 are “Human” and “Staphylococcus”). Then, the input files can be copied to their corresponding input folders. After the input files have been provided, READemption 2.0 automatically manages the input and output of all following subcommands. The next step is aligning the reads to the combined reference sequences of all species, using the subcommand ‘align’. It has been shown that aligning read pools, containing sequences from multiple species, to combined reference sequences instead of aligning the reads subsequently to each species reference genome avoids introducing mapping *bias**(Espindula et al., 2020***), which makes this combined approach READemption’s method of choice when analyzing Dual RNA-seq data. The mapping statistics generated by the ‘align’ subcommand were updated to include mapping statistics by species. These include counts for reads that align to a single species and reads that cross-align to multiple species (Figure 2C). Another new feature of the ‘align’ subcommand is the possibility to merge the two aligned reads of a read pair and build template fragments when analyzing paired-end data. The derived fragments are stored in a BAM file as single-end alignments and can be used for further analysis instead of the BAM files that include the un-merged paired-end reads. After running the ‘align’ subcommand the user can perform the subcommands ‘coverage’ or ‘gene quanti’ followed by ‘deseq’. The subcommand ‘coverage’ creates strand specific coverage files in wiggle format, containing nucleotide-wise read counts for the genomic positions of the reference sequences. The counts are provided with and without normalization and can be viewed in a genome browser for further inspection. The subcommand ‘gene quanti’ calculates the number of reads overlapping with each feature listed in the annotation files. The feature types to be used for the calculation can be specified by the user. The results are presented as raw counts and normalized counts, including transcripts per million (TPM, ***Wagner et al. (2012)***), reads per kilobase million (RPKM, ***Mortazavi et al. (2008)***) and normalized by the total number of aligned reads of the given library (TNOAR). After the gene quantification has been completed the subcommand ‘deseq’ can be used to perform differential gene expression analysis using the R package DESeq2 (***Love et al., 2014***), which is integrated into READemption. The subcommand also produces PCA (principal component analysis) plots and heatmaps of the library compositions. Finally, READemption offers subcommands for further visualization. ‘Viz_align’generates histograms of the read length distributions, ‘viz_gene_quanti’ bar plots of the feature distribution and scatter plots comparing raw gene-wise quantification values for each library pair and ‘viz_deseq’ MA and Volcano plots.

**Figure 3.**
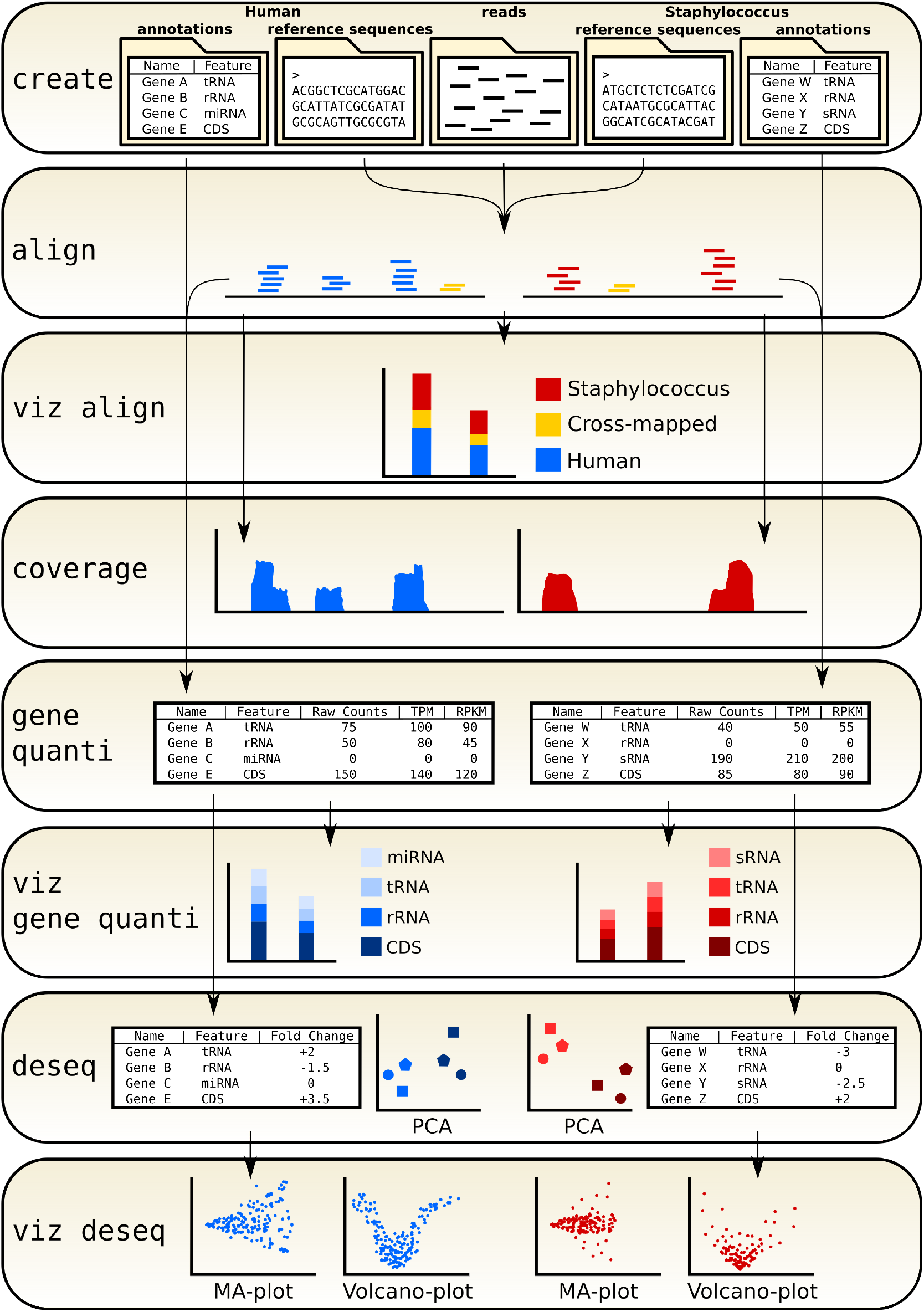
READemption 2.0 data and workflow overview of a Dual RNA-seq example analysis. Each subcommand is depicted as one box. The arrows indicate the data flow of the input and output

### Species cross-mapped reads and normalization

During the alignment the majority of reads can be unambigously assigned to their species. These species exclusive reads are then used for down-stream analysis of the corresponding species. Each subcommand that is being called after the initial alignment produces independent results for the different species. E.g. in a Dual RNA-seq experiment containing human cells and *Staphylococcus aureus* cells, READemption 2.0 creates coverage files once for the human genome and once for the bacterial genome, while only taking reads into account that map to the respective species (Figure 3: ‘Coverage’-box). However, a typical Dual or Multi RNA-seq experiment contains a small fraction of reads that map equally well to two or more species. These species cross-mapped reads pose a problem, since discarding them causes information loss while keeping them results in potential false positives. Although species cross-mapped reads are usually discarded, there is no gold standard of how to handle them (***Espindula et al., 2020***). To give users full control over cross-mapped reads and normalization, we added options to include or exclude cross-mapped reads for the nucleotide-wise counts of ‘coverage’ and the gene-wise counts of ‘gene quanti’, as well as including or excluding them in the values used for normalization for these subcommands. As cross-mapped reads are usually discarded, the default setting of READemption 2.0 is to exclude cross-mapped reads for both individual counts and normalization. READemption 2.0 provides three different ways for DESeq2’s size factor calculation that is used for normalizing read counts over different libraries. The project-wise approach takes all feature counts of a species of all libraries into account when comparing conditions with each other, the species-wise approach uses only the libraries of the given species, and the comparison-wise approach only the libraries of the two conditions that are currently compared. We chose the species-wise approach as default setting, since the ‘deseq’ subcommand also generates PCA plots based on the libraries used for size factor estimation and usually the first quality control step of differential gene expression analysis is confirming via PCA, whether the libraries of the same condition cluster together.

### Fragment building

Some manufacturers, e.g. Illumina or Applied Biosystems offer RNA-seq protocols that generate paired-end reads, where each cDNA template fragment is sequenced from both ends, resulting in a read pair. After the alignment the mapped pairs can be used to derive the genomic start and end position of the template they originate from. READemption 2.0 uses the alignment files (BAM files) of the initial alignment to generate template fragments from paired-end reads and writes them to a new BAM file containing the template fragments represented as single-end reads. Building these fragments is the default option, but can also be turned off to use the individual reads of a pair as input for the down-stream analysis.

## Discussion

The growing number of research articles using Dual RNA-seq (***PubMed, 2022b***) shows the need in the scientific community for a tool that can conveniently analyze the data of such experiments. We present READemption 2.0, the first tool that can handle RNA-seq data of any number of species and any domain of life. Other tools that already exist and are suitable for the analysis of Multispecies RNA-seq either provide only basic alignment functionalities or can only handle a maximum of two different species. FastQ-screen (https://stevenwingett.github.io/FastQ-Screen/) generates read files that contain information about to which species a read could be aligned. These reads can be filtered and used as input for other third-party tools. However, FastQ-screen does not provide coverage-file creation, gene-wise quantification or differential gene expression analysis. The nf-core/dualrnaseq pipeline (https://nf-co.re/dualrnaseq) is able to perform read alignment and gene-wise quantification, but lacks the ability to analyze more than two species. Although READemption 2 has been developed with the intent to analyze Dual or Multi RNA-seq data of interacting species, its application in other areas of RNA-seq is conceivable. E.g. metatranscriptome analysis similar to ***Krohn-Molt et al. (2017*)** could be conveniently analyzed with READemption 2.0. During the development of READemption 2.0 we focused on easy accessibility, to ensure that researchers can run analyses with little prior knowledge of bioinformatics. We did this by choosing default parameters that are most common for the analysis of either Dual and Multi RNA-seq or conventional RNA-seq and by providing comprehensive tutorials, explanations and solutions for convenient installation of the tool and all its dependencies. However, parameters can be changed in different ways (e.g. different normalization approaches, use of single reads or fragment building for paired-end reads etc.) to meet the users’ specific needs. This principal is called “convention of configuration” and has been applied to the default settings of all subcommands.

## Methods and Materials

READemption 2.0 is written in Python and the source code is freely available under the ISC license on GitHub (https://github.com/foerstner-lab/READemption). The short read mapper segemehl (***Hoffmann et al., 2009***) and the R package DESeq2 (***Love et al., 2014***) are integrated into READemption 2.0. Software unit and system tests were created to guarantee READemption 2.0 runs as intended. The tests cover 85 % of the code, including all core functions. READemption can be installed via Conda (https://anaconda.org/Till_Sauerwein/reademption), PyPi (https://pypi.org/project/READemption/) or using a pre-installed Docker image (https://hub.docker.com/r/tillsauerwein/reademption). READemption 2.0’s Documentation website (https://reademption.readthedocs.io/en/latest/index.html) hosts detailed descriptions of the subcommands, information about fragment building for paired-end reads, installation instructions and tutorials for beginners. The tutorials offer step-by-step instructions, input data and executable code to perform single or dual RNA-seq example analyses.

## Acknowledgments

We thank Silvia Di Giorgio and Muhammad Elhossary for testing READemption 2.0 and giving us feedback. This work was funded by the Deutsche Forschungsgemeinschaft (DFG, German Research Foundation) – Project number 460129525 (NFDI4Microbiota – National Research Data Infrastructure for Microbiota Research) and was supported by the IZKFatthe University of Würzburg (project Z-6).

